# Pseudocontact shift NMR data obtained from a non-canonical amino acid-linked lanthanide tag improves integral membrane protein structure prediction

**DOI:** 10.1101/2022.09.14.507970

**Authors:** Kaitlyn V. Ledwitch, Georg Künze, Katherine Larochelle, Elleansar Okwei, Lisa Pankewitz, Soumya Ganguly, Heather L. Darling, Irene Coin, Jens Meiler

## Abstract

A single experimental method alone often fails to provide the resolution, accuracy, and coverage needed to model integral membrane proteins (IMPs). Integrating computation with experimental data is a powerful approach to supplement missing structural information with atomic detail. We combine RosettaNMR with experimentally-derived paramagnetic NMR restraints to guide membrane protein structure prediction. We demonstrate this approach using the disulfide bond formation protein B (DsbB), an α-helical IMP. We attached a cyclen-based paramagnetic lanthanide tag to an engineered noncanonical amino acid (ncAA) using a copper-catalyzed azide-alkyne cycloaddition (CuAAC) click chemistry reaction. Using this tagging strategy, we collected 203 backbone H^N^ pseudocontact shifts (PCSs) for three different labeling sites and used these as input to guide *de novo* membrane protein structure prediction protocols in Rosetta. We find that this sparse PCS dataset combined with 44 long-range NOEs as restraints in our calculations improves structure prediction of DsbB by enhancements in model accuracy, sampling, and scoring. The most accurate DsbB models generated in this case gave Cα-RMSD values over the transmembrane region of 2.11 Å (best-RMSD) and 3.23 Å (best-scoring).

## Introduction

Integral Membrane Proteins (IMPs) are a complex class of proteins, including large macromolecular machines that perform biological tasks essential to human health. Access to atomic-level information on the structure and dynamics of these biomolecules is key to understanding their functions and mechanisms. Conventional biophysical techniques in structural biology like nuclear magnetic resonance (NMR), X-ray crystallography, and cryo-electron microscopy struggle to obtain experimental data that unambiguously define structural details for IMPs. For this reason, it is often necessary to combine sparse experimental data as restraints with computational modeling algorithms to supplement missing information with atomic detail.

NMR-derived restraints interrogate individual atoms of a biomolecule and are one of the most useful sources of structural information. A complete dataset of chemical shifts (CSs) and nuclear Overhauser effects (NOEs) can be used as restraints to determine unique structural models for small soluble proteins by NMR. NOEs represent the transfer of spin polarization between two sets of active populations of spin-active nuclei and can be measured at a maximum proton-proton distance of ~5 Å through a cross-relaxation phenomenon. This type of NOE-based geometric information between spins is important in NMR structural calculations. However, for IMP systems the large size leads to spectroscopic challenges due to slower tumbling of the protein in solution and broadening of line widths, complicating the acquisition of long-range NOEs. The average transmembrane helix-helix distance is typically in the range of 10 Å and 11 Å, although notable exceptions exist^1^. Thus, NOEs between backbone hydrogen atoms of different transmembrane helices are typically absent. This sparseness of data is insufficient for defining the IMP fold. Sidechain hydrogen atoms get close enough for proton-proton NOE measurements, but their assignment is difficult. Even with selective ^1^H-labeling of isoleucine, valine, and leucine methyl groups^2–12^, long-range NOEs remain sparse.

Advancements in technology such as the transverse relaxation Overhauser spectroscopy (TROSY) experiment^10,13–19^, sample perdeuteration^20,21^, and non-uniform sampling^22–25^ have pushed structure determination to protein sizes ranging from to 50–100 kDa. These technical developments have contributed to a range of detergent-solubilized NMR structural models deposited in the Protein Data Bank (PDB) including the outer mitochondrial voltage-dependent anion channel VDAC-1^26,27^, the archaeal phototaxis receptor sensory rhodopsin II pSRII^28^, and the bacterial inner IMP DsbB^29^. The recent emergence of small covalently-circularized nanodiscs has made it possible to determine structural models for the β-barrel proteins OmpX, OmpA, and Ail in a lipid environment^30–32^. This was also done for the neurotensin receptor NTR1 using a selectively modified sequence to stabilize the protein in a conformation amenable to NMR studies^31^. Even with these advances, it still remains a practical challenge for structure determination of α-helical IMPs using traditional NMR restraints, especially those containing more than seven transmembrane helices^30^.

The introduction of a paramagnetic tag is an alternative strategy to obtain long-range NMR restraints for the 3D structural modeling of α-helical IMPs^33^. The paramagnetic center leads to an interaction between the unpaired electron and the nuclear spins of the protein generating NMR observables such as pseudocontact shifts (PCSs), residual dipolar couplings (RDCs), and paramagnetic relaxation enhancements (PREs). PREs can be detected in any paramagnetic system and can be used to extract distances. PCSs and RDCs depend on anisotropic magnetic susceptibility of the paramagnetic ion (e.g., lanthanide ion), giving rise to a non-vanishing magnetic susceptibility anisotropy (Δχ)-tensor and partial alignment of the protein in the magnetic field. PCSs arise from a through-space interaction between the unpaired electron and the nucleus and are particularly useful as restraints because they combine both distance and angular information. PCSs are measured as the difference in chemical shift between a paramagnetic and diamagnetic sample and stand out over the other types of paramagnetic effects due to the ease of measurement. The acquisition and use of PCSs is a powerful source of spatial information and can be exploited in various ways to probe the structure, function, and dynamics of biomacromolecules.

The conventional method is to introduce a paramagnetic lanthanide-based spin-label to the protein of interest through a site-specific cysteine residue via a disulfide linkage. This requires the removal of all native cysteines and the site-specific reintroduction of single cysteine mutants. However, endogenous cysteine residues are inherently important to the overall three-dimensional fold, stability, and function of a protein and in some cases, cannot be removed. For example, G-protein coupled receptors (GPCRs) have disulfide bridges that stabilize the extracellular loops and are key to biological function. The disulfide bond formation protein B (DsbB) was the first NMR-derived model for a polytopic IMP and plays a key role in disulfide bond formation in *E. coli*. This structural model represents the interloop disulfide bond intermediate and was solved in part using PREs derived from nine single cysteine mutants^29^. Interestingly, Zhou *et al*. used a construct for these paramagnetic measurements that retained an interloop disulfide bond between two native cysteine residues at positions C41 and C130, allowing two of the six native DsbB cysteines to be kept intact^29^.

An orthogonal strategy to circumvent the need to remove all native cysteines is the use of genetically encoded noncanonical amino acids (ncAAs) that either have a paramagnetic center or a reactive group for the attachment of a spin-label. The site-specific incorporation of the ncAA *p*-azido-ʟ-phenylalanine (pAzF) and ligation with alkyne-bearing lanthanide tags (C3 and C4) via a copper-catalyzed azide-alkyne cycloaddition (CuAAC) click chemistry reaction was first reported by Loh *et al*. for several soluble proteins including ubiquitin, sortase A, and T4-lysozyme^34–36^. The synthesis of the 1,4,7,10-tetraazacyclododecane (cyclen)-based C3 and C4 lanthanide tags are based on the cysteine-compatible C1 and C2 tags^37,38^. The C1, C3, and C4 lanthanide tags share the same phenylethylamide pendant chirality whereas the C2 tag is the same molecule of C1, just the opposite enantiomer. The C3 tag has a shorter tether linking the paramagnetic lanthanide to the protein backbone compared to the C4 tag and for the soluble test cases mentioned above, the C3 tag generated PCS datasets with better quality (Q)-factors. Few examples in the literature have used these tags for PCS-guided structure determination of IMPs. In one example, 737 PCS-derived restraints from a C2 tag were used in combination with a limited set of NOEs to determine the global fold of the backbone for the 7 transmembrane ⍺-helical microbial receptor pSRII^39^. This study showed that the NOEs alone were insufficient to correctly determine the three-dimensional (3D) fold, emphasizing the utility of PCSs to provide complementary information.

Here, we use detergent-solubilized DsbB as a model IMP system to attach the C3 lanthanide spin-label to an engineered pAzF ncAA using a CuAAC click chemistry reaction. To our knowledge, this is the first test case demonstrating the utility of this labeling scheme in generating PCSs as input for structural modeling of IMPs. We use DsbB as a model IMP because 1) cysteine residues (e.g., C41–C130) are catalytically critical in the DsbB reaction cycle, 2) using a ncAA will avoid the need to introduce a non-native cysteine residue for generating long-range paramagnetic data, and 3) the deposited NMR structural ensemble allows us to evaluate the performance of traditional NMR data against the utility of sparse PCS restraints for calculating the three-dimensional fold of DsbB. To generate paramagnetic NMR data, the C3 tag was loaded with the lanthanide ion ytterbium (Yb^3+^) and we collected 203 H^N^ backbone PCSs from three different pAzF tagging sites (F32, V72, and Y97). These PCSs were used as input to test the performance of sparse PCS data in guiding *de novo* folding of DsbB in RosettaNMR—a unified framework for using traditional NMR data with paramagnetic NMR data for a number of structural modeling tasks^40^. We compared these results to the performance of the traditional NMR restraints (chemical shifts, NOEs) used to calculate the NMR-derived structural ensemble of DsbB (PDB ID: 2K73). We demonstrate that including this sparse PCS dataset as restraints improves structure prediction of DsbB by enhancements in model accuracy, sampling, and scoring. This PCS-driven integrated modeling framework is equally applicable to other IMP systems, especially those where the removal of native cysteine residues complicates biophysical experiments or the reintroduction of single cysteines leads to the formation of undesired dimeric states.

## Results and Discussion

### Near-quantitative CuAAC ligation yields for DsbB-pAzF mutants are achieved by decreasing the total protein induction time

Figure 1 shows a schematic highlighting the location of the selected DsbB sites for pAzF insertion, the C3-lanthanide tag, and the formation of the triazole ring between the pAzF azide group and the C3 alkyne group following the CuAAC reaction. Residues F32, V72, and Y97 (Figure 1, grey spheres) were selected as sites for pAzF incorporation based on the criteria that they are solvent-exposed and either an aromatic residue or one of the nine single cysteine sites used for the deposited NMR structure calculation. Analogous to the global positioning system, three sites were selected for spin-labeling and are distributed over the protein space for determining secondary structural elements within the protein fold by triangulation. A range of reaction components were tested to determine optimal conditions for increasing the CuAAC ligation yields and include the addition of the Cu(I) stabilizing ligand BTTAA, glycerol, and aminoguanidine^36^. Near-quantitative yields for the CuAAC reaction with the C3-tag were reported for the soluble protein sortase A upon removal of the His_6_ tag used for purification^36^. We performed the CuAAC ligation reactions for DsbB and the C3-tag based on these optimized conditions with the exception that the C-terminal His_6_ tag was not removed.

**Figure 1.**
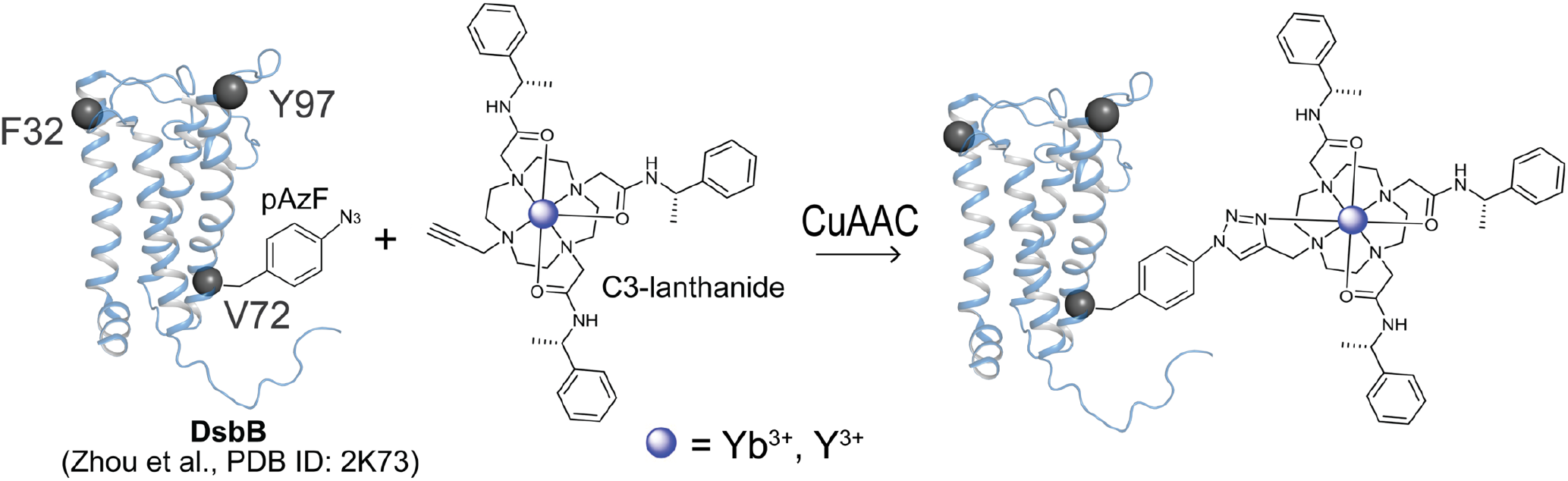
Schematic showing the copper-catalyzed azide-alkyne cycloaddition (CuAAC) click chemistry reaction. We engineered three different pAzF DsbB mutants for attaching a paramagnetic cyclen-based C3-lanthanide tag at sites F32, V72, and Y97 (grey spheres) using a copper-catalyzed click chemistry reaction. The CuAAC reaction yields an azide-alkyne cycloaddition product incorporating the paramagnetic or diamagnetic C3-lanthanide ion tag at a site-specific pAzF location in the protein system.

It is also well recognized that azide groups are sensitive to reduction during *E. coli* expression and purification, decreasing the C3-tag ligation reaction efficiency^41^. We tested a range of induction times for expressing the V72 DsbB-pAzF mutants to determine the time-dependent reduction of pAzF on azide viability for the covalent attachment of the C3 spin-label. DsbB-pAzF mutants were induced for three, five, and eight hours of total expression time. We calculated the percentage of successfully ligated protein by measuring the relative cross-peak intensities between the ligated protein and the corresponding peak for the unligated protein. The NMR signal from the ligated protein is the shifted peak due to the PCS effect and the NMR signal from the unligated protein is the unshifted peak in the paramagnetic spectrum that corresponds to the diamagnetic reference. The ligation efficiencies increased with respect to shorter induction times and were 99%, 65%, and 26%, respectively. In this case, we found that complete ligation reaction efficiencies had a negative impact on the spectral quality and measurement of PCSs. We hypothesize that this is most likely the result of a strong PRE effect which causes the broadening of peaks in the proximity of the paramagnetic tag past the level of detection. In the case of DsbB, we found that partial ligation yields were beneficial because it allowed for easier assignment of paramagnetic and diamagnetic peaks in the NMR spectrum. Because partial ligation yields do not interfere with the determination of accurate Δχ tensors, we used an induction time of five hours to maintain partial ligation yields to balance accurate assignment of resonances and sizeable PCS effects.

### Solution NMR spectra of C3-labeled DsbB at position Y97pAzF generates the largest and most pronounced PCS dataset

Figure 2 shows a representative ^1^H-^15^N TROSY-HSQC NMR spectrum for paramagnetically-labeled (blue) DsbB at position Y97pAzF superimposed with the diamagnetic reference spectrum (red). Zoomed in regions of cross-peaks show the most pronounced PCSs. The C3-tag was loaded with the lanthanide ion ytterbium (Yb^3+^) and yttrium (Y^3+^) to record the paramagnetic and diamagnetic spectrums, respectively. The ^1^H-^15^N 2D NMR spectral fingerprint for all pAzF mutants were directly comparable to the published ^1^H-^15^N 2D NMR spectrum for DsbB^29^. We transferred the published backbone chemical shifts from the BMRB (BMRB: 15966) to assign each PCS for each DsbB-pAzF mutant. We were able to unambiguously assign a total of 99 PCSs in the range of −0.55 ppm to +0.17 ppm for Y97pAzF. This represented the largest and most pronounced PCS dataset compared to the dataset for the other tagging sites. For the F32pAzF and V72pAzF tagging sites, we measured 60 PCSs and 54 PCSs with magnitudes in the range of −0.14 ppm to +0.36 ppm and −0.12 ppm to +0.19 ppm, respectively. The ^1^H-^15^N TROSY-HSQC NMR spectra for these two additional datasets are shown in Figure S1. All observed ^1^H^N^ PCS values for each DsbB C3 tagging site are reported in Table S1.

**Figure 2.**
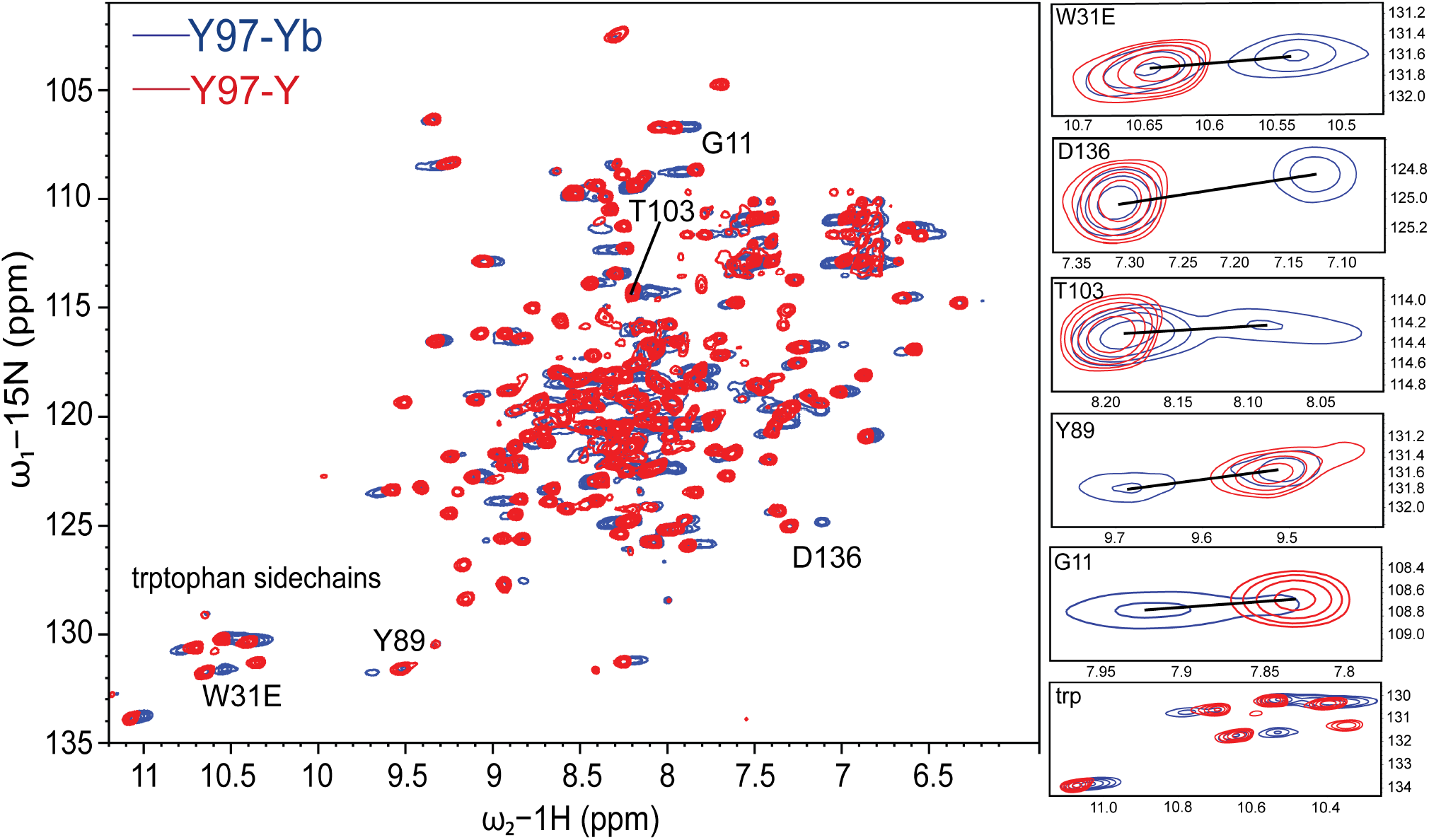
Representative ^1^H-^15^N TROSY-HSQC NMR spectra for para- and diamagnetically-labeled DsbB at position Y97pAzF. The ^1^H-^15^N TROSY-HSQC spectrum for the paramagnetic C3-Yb^3+^ sample (blue) is superimposed with the spectrum of the diamagnetic C3-Y^3+^ sample (red) to showcase the paramagnetic PCS effects for pAzF-tagged DsbB at amino acid position Y97. NMR data was collected at 40°C and a proton resonance frequency of 900 MHz. Zoomed in regions (right) highlight DsbB residues that showed the most pronounced PCS effects.

We also incorporated pAzF at sites A14 and Y46 and attempted CuAAC reactions with the C3-Yb^3+^ spin label for NMR measurements. The A14 site was one of the nine sites used for cysteine labeling in generating PREs for the deposited NMR structural ensemble. We selected Y46 because this residue not only faces the solvent, but was further motivated by the rationale that replacing an aromatic residue with a ncAA aromatic with similar physiochemical properties would introduce minimal perturbations to the native structure. Unfortunately, these selected sites for tagging either resulted in poor spectral quality or the observations of PCSs effects could only be assigned for a few resonances. For the A14pAzF tagging site, the ligation efficiency at a total protein induction time of five hours was 34%, which was substantially lower than the expected range. A14 is localized to transmembrane helix one (TM1) and based off published EPR studies that report the depth of nitroxide insertion in the membrane bilayer, A14 flanks the lipid bilayer plane in close proximity to helix one (H1) off of the N-terminus^29^. One explanation is that this position can accommodate a small nitroxide spin label and not a bulky C3 tag due to added steric hindrance from the insertion of a larger amino acid. We had success with the insertion of pAzF at position Y46 but interestingly, the attempted isolation of the protein resulted in an uncharacteristic shift in the color of the purified protein solution from yellow to purple and subsequent tagging reactions were not attempted.

### Δχ-tensors generated for the C3-Yb^3+^ tag give reasonable Q-factors

The Δχ tensors for C3-Yb^3+^ labeled DsbB at positions F32pAzF, V72pAzF and Y97pAzF were each fit to the first conformer of the DsbB NMR structural ensemble (PDB ID: 2K73) and are summarized in Table 1. Figure 3 shows the correlation between the experimental and calculated ^1^H^N^ PCS values (Figure 3, left). Visualization of the anisotropy parameters induced by the paramagnetic Yb^3+^ ion is shown in the form of PCS isosurfaces for each mutant (Figure 3, right). The fit of back-calculated versus experimental PCSs gave reasonable Q-factors of 0.365, 0.309, and 0.191 for C3-Yb labeled F32pAzF, V72pAzF and Y97pAzF, respectively.

**Table 1:** Δχ Tensor parameters for DsbB - pAzF mutants (F32, V72, Y97) tagged with C3-Yb^3+^.

| C3—Yb<sup>3+</sup> | $\Delta\chi_{ax}$<br>(10<sup>-32</sup> m<sup>3</sup>) | $\Delta\chi_{rh}$<br>(10<sup>-32</sup> m<sup>3</sup>) | Q | x<br>(Å) | y<br>(Å) | z<br>(Å) | $\alpha$<br>(°) | $\beta$<br>(°) | $\gamma$<br>(°) |
| --- | --- | --- | --- | --- | --- | --- | --- | --- | --- |
| F32 | 1.126 | 0.493 | 0.365 | -8.170 | 12.992 | 18.188 | 42.808 | 163.953 | 151.035 |
| V72 | 4.283 | 0.870 | 0.309 | -4.774 | -15.998 | -8.484 | 92.986 | 30.262 | 155.521 |
| Y97 | 12.470 | 3.056 | 0.191 | 18.553 | 12.024 | 0.224 | 84.928 | 111.407 | 78.248 |

<sup>a</sup> The Euler angles ( $\alpha$ , $\beta$ , $\gamma$ ) for the PCS tensors are in zyz convention in units of degrees.
<sup>b</sup> The metal coordinates (xyz) and tensor parameters are reported relative to the first conformer of the NMR structure of DsbB (PDB ID: 2K73).
<sup>c</sup> Quality (Q) factors were calculated as the root-mean-square deviation between experimental and back-calculated PCSs divided by the root-mean-square of the experimental PCSs.

**Figure 3.**
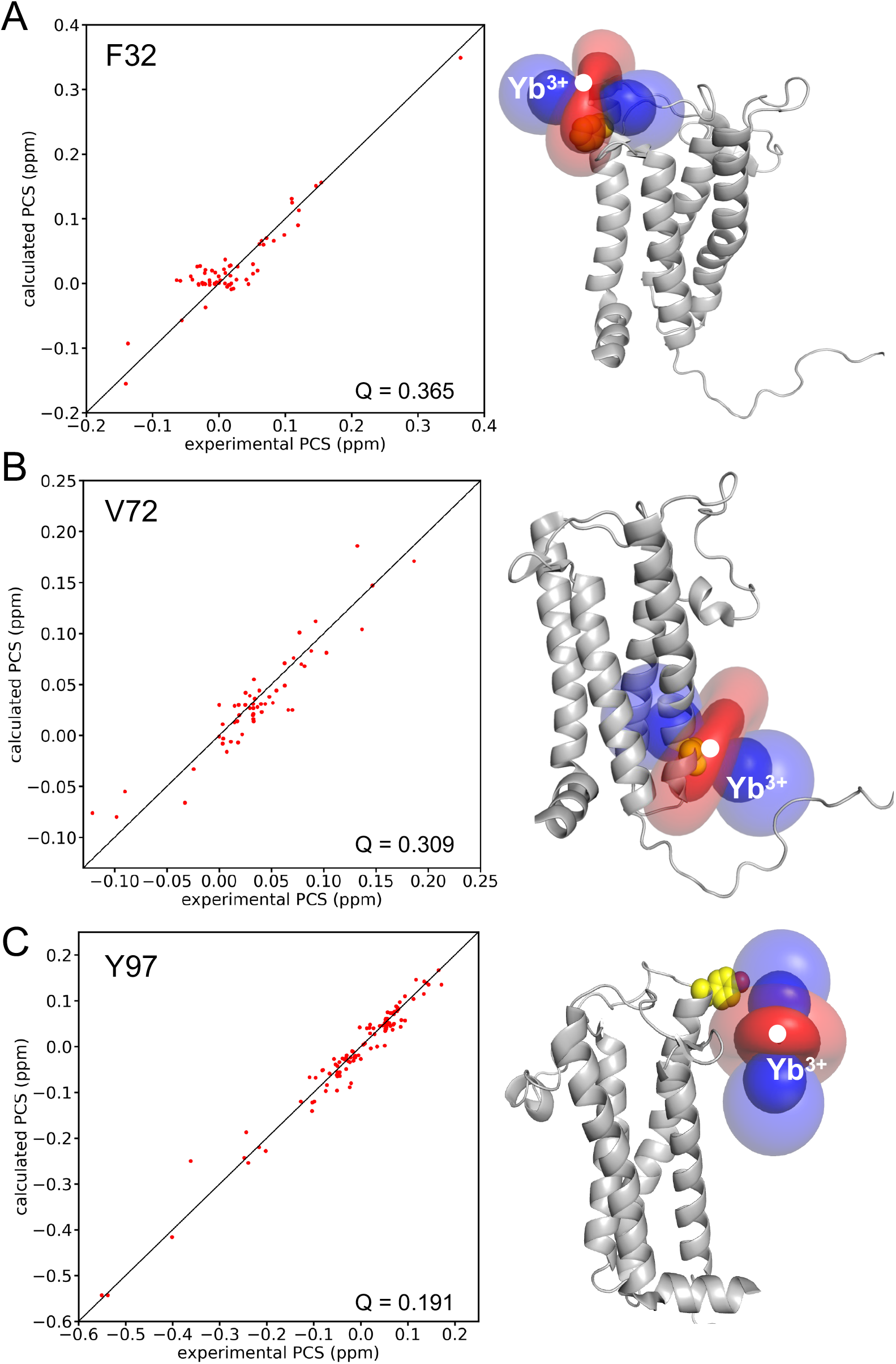
Magnetic susceptibility anisotropy (Δχ) tensor fits to the first conformer of the NMR structural ensemble for DsbB (PDB ID: 2K73). Correlation between experimental and calculated ^1^H^N^ PCS values (left), and the isosurface representation of the PCS tensor (right) for the paramagnetic Yb^3+^ ion coordinated through the C3-labeled pAzF DsbB mutant (**A**) for F32pAzF - DsbB (**B**) for V72pAzF - DsbB (**C**) and for Y97pAzF - DsbB.

The Y97pAzF labeling site produced the most pronounced PCS dataset for DsbB in terms of both PCS quantity and PCS size compared to F32pAzF and V72pAzF. The magnitude of the Δχ tensor for Y97 is considerably larger than the tensors determined for the other two C3 labeling sites (Table 1). Loh *et al*. determined the Δχ tensors for C3-labeled ubiquitin at two different pAzF positions and showed that the tensor for T66pAzF was 2-fold larger than G18pAzF^42^. They concluded this was potentially due to interactions between the tag and the protein, increasing the rigidity of the tag and therefore the magnitude of the Δχ tensor. In the case for DsbB, residue Y97 is localized on the cusp of periplasmic loop two (PL2) and determined to be outside of the lipid bilayer plane^42^. It seems unlikely that the C3 tag at this position would make interactions with DsbB. Instead, we take into consideration that unlike the other positions, this amino acid would be positioned directly above the detergent micelle. We speculate here that the magnitude of the Δχ tensor for Y97pAzF and quality of the PCS dataset is the result of the proximity of the detergent micelle to Y97 inadvertently increasing the rigidity of the tag. In retrospect, we also argue that the lack of the detergent micelle around Y97 is advantageous to the rigidity of the tag compared to spin labeling sites that would be surrounded by detergent.

### PCSs facilitate *de novo* structure prediction of DsbB by improvements in model sampling and scoring

We tested the performance of PCSs for guiding IMP structure prediction by calculating the structure of DsbB with Rosetta using our experimental PCS data. A total of 203 backbone H^N^ PCSs in the range from −0.55 ppm to +0.36 ppm could be used. This included 60 PCSs for tagging site F32, 54 PCSs for tagging site V72, and 99 PCSs for tagging site Y97 (Table S2). Overall, this represented a sparse PCS dataset and a difficult test case for protein NMR structure determination.

We employed the Rosetta fragment assembly protocol for structure calculation. PCSs were used at all four stages of the protocol to score trial structures that are created along the folding trajectory and guide model sampling. Two different fragment libraries were tested, one created with sequence-based secondary structure information for DsbB and another created with chemical shift (CS) data for DsbB. The backbone CS data could be obtained from the published NMR assignment of DsbB (BMRB: 15966)^29^ and were used to derive φ/ψ angle and secondary structure predictions as input for the Rosetta fragment picker application. Structure calculation performance was evaluated by model accuracy in terms of the Cα-atom RMSD calculated over all residues in DsbB or only residues in transmembrane segments (RMSD_TM) of DsbB. In addition, we compared the performance of structure calculations with PCSs to that obtained with a sparse dataset of 40 long-range NOEs (“lr_NOE”) as well as a larger dataset including 500 short-range and long-range NOEs (“all_NOE”). The NOE dataset was also obtained from the published NMR assignment of DsbB (BMRB: 15966)^29^. Figure 4 summarizes the Cα-RMSD and Cα-RMSD_TM values of the best-scoring and best-RMSD models of DsbB generated with PCSs and other types of NMR data. Figures S2 and S3 depict the folding energy landscape of the corresponding structure calculations by showing their model score-versus-RMSD plots.

**Figure 4.**
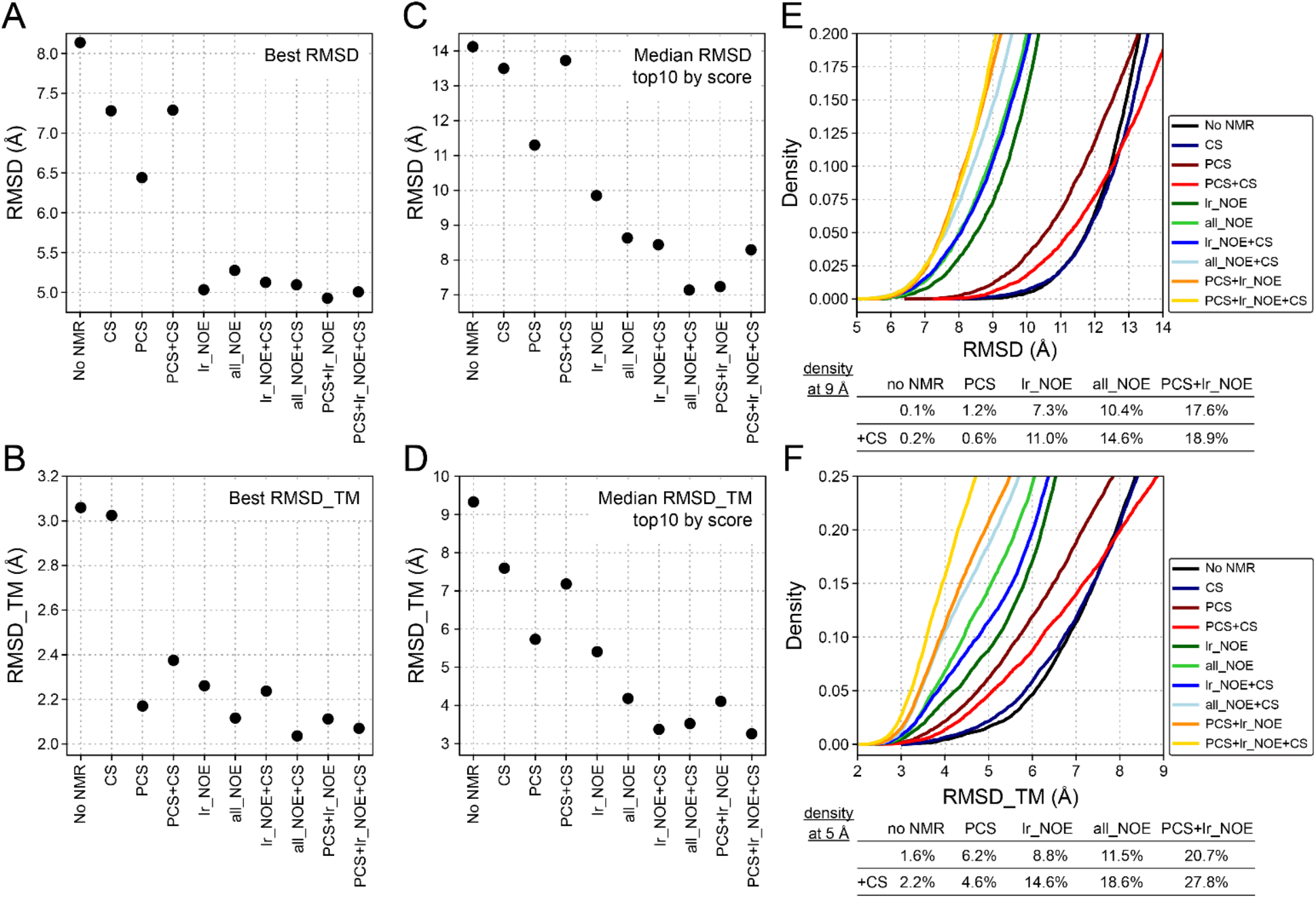
Summary of RMSD values of DsbB models predicted with Rosetta using PCSs and other types of NMR data. (**A**) Cα-RMSD values over all amino acid residues of the lowest-RMSD DsbB models predicted in the absence (no NMR) or presence of different types of NMR data. CS – chemical shifts, PCS – pseudocontact shifts, lr_NOE – long-range NOEs, all_NOEs – all short-range and long-range NOEs available for DsbB (BMRB: 15966). (**B**) Cα-RMSD values over all residues in transmembrane regions of the lowest-RMSD_TM models of DsbB predicted with Rosetta and different types of NMR data. (**C**) Median Cα-RMSD and (**D**) median Cα-RMSD_TM values of the ten best scoring DsbB models that were predicted and scored using either the Rosetta score3 energy function or a combination of the score3 energy and one of different types of NMR restraints. (**E**) Density of DsbB models as a function of model Cα-RMSD value in structure calculations with PCSs and other types of NMR data. (**F**) Density of DsbB models as a function of their Cα-RMSD_TM value.

Using PCSs improved the model accuracy compared to the case when no NMR data or only CS-derived fragments were used. The best-scoring and best-RMSD models of DsbB created with PCSs had Cα-RMSD_TM values of 5.73 Å and 2.18 Å, respectively, whereas without NMR data the RMSD_TM values were 9.57 Å and 3.06 Å (Figure 4A+B). Using the sparse set of long-range NOEs further improved the RMSD_TM for the best-scoring and best-RMSD models to 3.53 Å and 2.26 Å, respectively. A comparable improvement was observed with the larger NOE dataset (all_NOE), yielding best-scoring and best-RMSD models with RMSD_TM values of 4.09 Å and 2.23 Å, respectively. The most accurate DsbB models were created when PCSs and long-range NOEs were combined. RMSD_TM values of 3.23 Å and 2.11 Å were reached for the best-scoring and best-RMSD models in this case. This order of RMSD improvement is also observed when looking at the median all-residue RMSD (Figure 4C) and median RMSD_TM (Figure 4D) values found for the ten best-scoring models of each structure calculation. The RMSD improves in the order: no NMR data > PCSs > lr_NOE/all_NOE > PCSs + lr_NOE. The gradual improvement of model accuracy becomes also evident by visual inspection of the Rosetta-generated models for DsbB. Figure 5 shows the best-scoring and best-RMSD models generated with no NMR data, PCSs, long-range NOEs, and combined usage of PCSs and long-range NOEs. In the best-scoring model created in the absence of PCSs and NOEs, the helix bundle packing is incorrect because the locations of TM helices 3 and 4 are swapped. By contrast, the best-scoring models created with PCSs or long-range NOEs have the correct helix bundle topology but some TM helices are unusually curved, which is a feature that is not observed in the experimental reference structure. In contrast, the best-scoring model created by combining PCSs and NOEs has the correct helix packing topology and helix shape, which is reflected by the very low Cα-atom RMSD value over the TM region of 2.62 Å.

**Figure 5:**
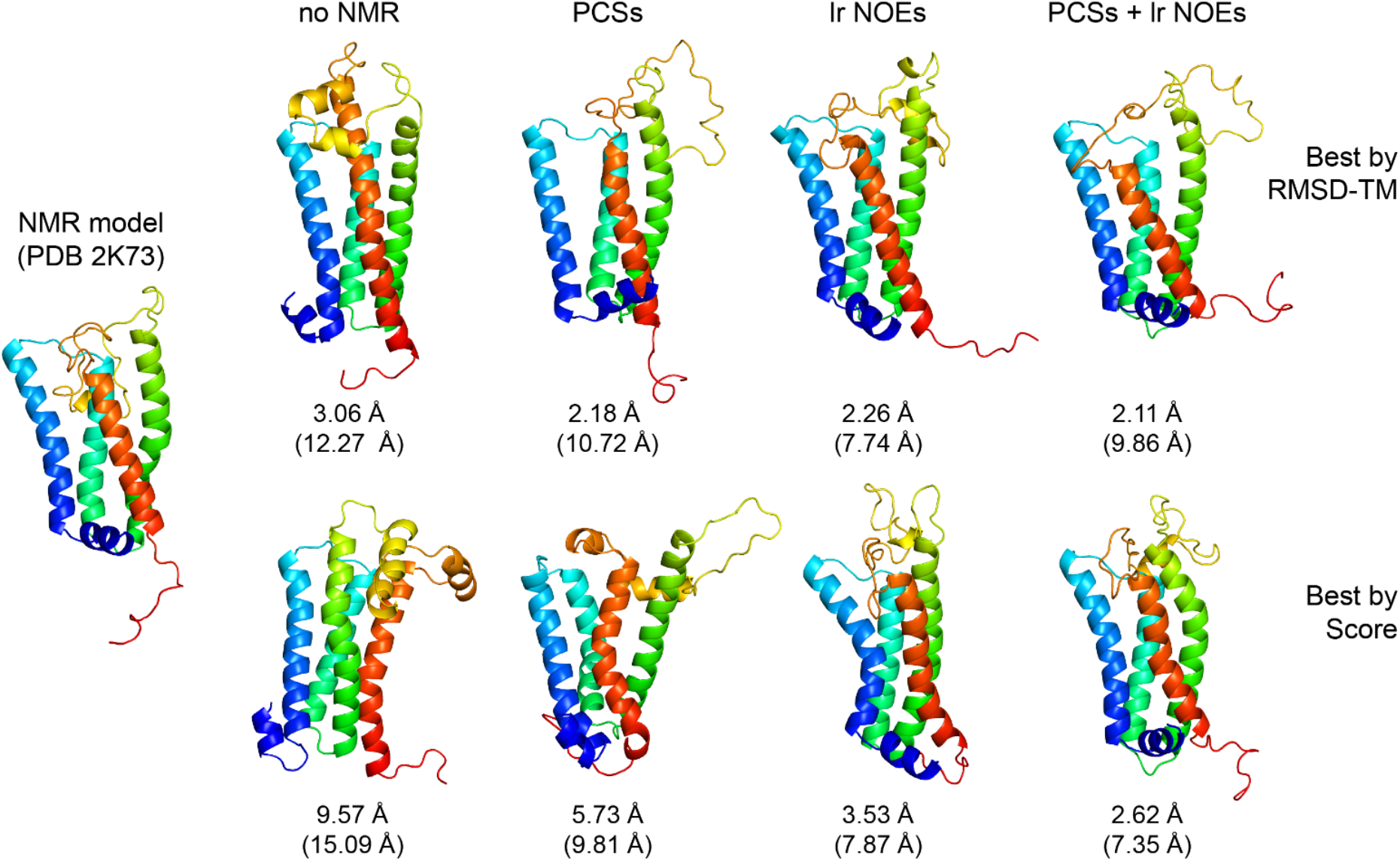
Cartoon representations of lowest-RMSD and lowest-scoring DsbB models predicted with Rosetta using PCS or NOE data. The NMR model of DsbB (PDB: 2K73) on the left side is compared with the best-RMSD_TM and best-scoring models of DsbB predicted in Rosetta structure calculations with either no NMR data, PCSs, long-range NOEs, or a combination of PCSs and long-range NOEs. The Cα-RMSD_TM (and all-residue Cα-RMSD) values are reported below each model.

Besides analyzing the best Rosetta models obtained for DsbB, we also investigated if PCSs improved the sampling of correctly-folded models in the fragment assembly protocol and the selection of more accurate models in the final scoring step. Figures 4E and 4F show the density of models that are better than a certain RMSD or RMSD_TM cutoff. Again, a gradual improvement in the fraction of most accurate models is observed in structure calculations with PCSs and NOEs compared to the calculation with no NMR data. For instance, the fraction of models with a RMSD_TM below 5 Å increased from 1.6% in the case with no NMR data to 6.2% when PCSs were used and to 8.8% when long-range NOEs were used (see Figure 4F). Combining PCSs and long-range NOEs led to an even more pronounced increase to 20.7%. A similar trend is seen when the all-residue RMSD of DsbB models is evaluated (Figure 4E). Using the CS-derived fragment library usually led to more low-RMSD models compared to calculations using the sequence-based fragment library for DsbB. Unexpectedly, CS-derived fragments could not improve model density in the PCS-guided structure calculation, but led to a small deterioration. However, a significant improvement was observed when CS-derived fragments were used together with PCSs and NOEs. 27.8% of models had a RMSD_TM smaller than 5 Å and 18.9% of models had an all-residue RMSD below 9 Å when PCSs, long-range NOEs, and CS data were used together.

Besides improvements in model sampling, we observed that PCSs had a favorable effect on model scoring and selection. When models generated in the absence of PCSs, either without any NMR data or with NOEs, were rescored with PCSs, it was possible to select a top-scoring model which had a lower RMSD value compared to the top-scoring model selected by considering only the plain Rosetta score or NOE restraint score. Figure S4 shows that the RMSD improvement by rescoring and selecting models with PCSs could be as large as 3 – 4 Å. Similarly, when models generated with PCSs were rescored with NOE restraints, lower RMSD models could be selected. This illustrates that PCSs and NOEs carry complementary structural information and that there is merit in combining both data for IMP structure calculations. In summary, we can conclude based on our test calculations for DsbB that PCSs improve structure prediction of this all-helical IMP by enhancements in model sampling and scoring. DsbB structure models with accuracies better than 3 Å in their TM region can be generated by using sparse PCS and NOE datasets.

## Conclusions

PCSs interrogate individual atoms of a biomolecule and offer a rich source of long-range structural information in the form of both distance and angular restraints. In addition to structure prediction, PCSs are sensitive to conformational rearrangements that typically escape traditional NOE measurements^43^. This type of information can be exploited as a tool to understand protein dynamics and when combined with ^15^N chemical exchange saturation transfer mapping (CEST)^44^, can be useful in identifying lowly-populated or short-lived conformational states. In the advent of AlphaFold, tools like this are key to the experimental validation of biophysically-relevant conformational states of proteins.

In summary, the present work demonstrates the utility of a pAzF-based C3 spin labeling scheme for an IMP that does not require cysteine mutagenesis for the introduction of a paramagnetic tag. We used this labeling strategy to generate a sparse PCS dataset from three different pAzF–DsbB sites and show that including this information as input for *de novo* structure calculations improves model accuracy, sampling, and scoring. The transfer of this labeling approach from soluble to IMPs expands paramagnetic NMR applications to larger systems and complexes, where traditional NMR datasets for structure calculations often fall short. Overall, the biocompatibility of pAzF and selectivity of the azido group for biorthogonal spin-labeling reactions offer a major advantage over conventional cysteine-based methods.

## Acknowledgments

This work was supported by NIH grants R01 GM080403, R01 HL122010, and R01 GM129261. The authors further acknowledge funding by the Deutsche Forschungsgemeinschaft (DFG) through SFB1423, project number 421152132. Jens Meiler is supported by a Humboldt Professorship of the Alexander von Humboldt Foundation. This work was conducted using the resources of the Biomolecular NMR facility and the Advanced Computing Center for Research and Education (ACCRE) at Vanderbilt University. We especially thank Prof. Markus Voehler for technical assistance with the NMR experiments. We thank Dr. Otting of the Australian National University for providing a test batch of the C3-based paramagnetic tag. All subsequent batches of the C3-based paramagnetic tag were synthesized in-house by Dr. Kwangho Kim of the Vanderbilt Institute of Chemical Biology, Molecular Design and Synthesis Center, Vanderbilt University, Nashville, TN 37232-0412.

## Author Contributions

Conceptualization, KVL, GK, and JM; Data curation, KVL, GK, KEL, EO, HD, LP, and SG; Formal analysis, KVL and GK; Funding acquisition, JM; Investigation, KVL, GK, and JM; Methodology, KVL, GK, and IC; Supervision, KVL and JM; Writing—original draft, KVL; Writing—review and editing, KVL, GK, and JM. All authors have read and agreed to the published version of the manuscript.

## Methods

### Synthesis of the C3-lanthanide tags

The cyclen-based C3 tags were synthesized as described^36^ and loaded with the paramagnetic ion ytterbium (Yb^3+^) or the diamagnetic ion yttrium (Y^3+^) for downstream click chemistry reactions and NMR experiments. Briefly, the synthesized 2,2’,2’’-(10-(prop-2-yn-1-yl)-1,4,7,10-tetraazacyclododecane-1,4,7-triyl)tris(N-((S)-1-phenylethyl)acetamide) C3 product (8 mg, 0.012 mmol) was dissolved in chloroform (1 mL) and basified with saturated NaHCO3 (2 mL). The basified C3 product was filtered through a phase separator with 10 mL of chloroform. The organic layer was removed *in vacuo* and the residue was dissolved in acetonitrile/water (0.5 mL/0.5 mL) followed by ytterbium(III) or yttrium(III) chloride (5 mg, 0.012 mmol) (Sigma Aldrich, ≥99.99%). The reaction mixture was heated at 85°C for 24 hours and the solvent was removed by lyophilization to afford the C3-lanthanide complex as a white solid product.

### Expression and purification of ^15^N-labeled DsbB-pAzF mutants

DsbB was expressed with a C-terminal His_6_-tag and an amber stop codon to replace residues Phe32, Val72, and Tyr97 with pAzF as described^34,36^. Protein genes containing the amber stop codon (UAG) were cloned in a pet22b vector to produce site-specific pAzF-DsbB mutants. C43 (DE3) *E. coli* cells were transformed with both the plasmid containing DsbB and the published pEVOL vector with the aminoacyl-tRNA synthetase machinery for pAzF insertion^45,46^. All transformations and cell cultures were grown at 37°C in the presence of 100 μg/mL ampicillin and 33 μg/mL chloramphenicol.

C43 (DE3)-pEVOL cells were freshly transformed with the DsbB-pAzF plasmid and grown on Luria-Bertani (LB) agar plates overnight at 37°C. Selected colonies were used to inoculate a 50 mL LB starter culture and grown overnight at 37°C. Cells from the starter culture were gently spun down at 4°C and resuspended in 1L of minimal (M9) media enriched with ^15^N-labeled ammonium chloride (^15^NH_4_Cl) and 0.02% arabinose. Cells were cultured at 37°C until the OD_600_ reached 0.6–0.8 and induced with 1 mM pAzF and 1 mM isopropyl-β-D-thiogalactopyranoside (IPTG). Cells were harvested by centrifugation at 6500 rpm at 4°C for 15 minutes within 3–5 hours after induction to preserve the integrity of the pAzF azide group. Cell pellets were stored at −80°C or immediately resuspended in lysis buffer for purification.

DsbB-pAzF mutants were purified as described previously^29^. Each 1L cell pellet was resuspended in 40 mL of Buffer A (50 mM Tris HCl, 300 mM NaCl, pH 8.0) containing 1 mM PMSF, one EDTA-free protease cocktail inhibitor tablet (Sigma-Aldrich), and 5 mM Mg Acetate. The cell lysis solution was rotated at 4°C for 30 minutes and then lysed by sonication for 10 minutes (60% amplitude with 5 seconds on/5 seconds off). The membrane fraction was pelleted by ultracentrifugation at 100,000xg for 1 hour. The pelleted membrane fraction was homogenized with Buffer A and 1% dodecylphosphocholine (DPC, FOS-CHOLINE-12, Anatrace) and was solubilized overnight at 4°C with gentle shaking. The insoluble fraction was removed by a subsequent centrifugation step 100,000xg for 30 minutes. Solubilized DsbB was isolated using Ni-NTA affinity chromatography using Buffer A and 0.1% DPC. The protein was eluted with 300 mM imidazole and concentrated in a 10 kDa MWCO Amicon device. The DsbB-pAzF samples were size excluded using a HiLoad® 16/600 Superdex® S200 preparative column (Cytiva) equilibrated with FPLC buffer (50 mM HEPES, 100 mM NaCl, 0.1% DPC, pH 7.5). The pooled fractions were concentrated in an Amicon centrifugal filter (10,000 MWCO) and analyzed by SDS-PAGE to confirm the purity of the sample was greater than 95%.

### Click chemistry reactions

The C3-lanthanide tags were prepared as a 10 mM stock in water by gently heating the stock solution to dissolve the tag and subsequently syringe filtered using a 0.2 μM filter. CuAAC reactions between the C3-lanthanide tags and the pAzF-incorporated protein were performed as described^36^ with the following modifications. All CuAAC reactions were performed in an anaerobic chamber under N_2_ atmosphere at room temperature to promote the reduction of Cu(II) to Cu(I) to facilitate efficient tagging. Reactions proceeded for 16 – 18 hours with gentle rotation. Reaction volumes were determined by the total amount of protein sample available and split in half to prepare matched samples for both the paramagnetic (C3—Yb^3+^) and diamagnetic (C3—Y^3+^) NMR samples, respectively. Final concentrations of the CuAAC reaction components included 0.1 mM pAzF-containing protein, 1 mM C3-tag, 2 mM of the copper-binding ligand BTTAA (2-(4-((bis[(1-tert-butyl-1H-1,2,3- triazol-4-yl)methyl]-amino)methyl)-1H-1,2,3-triazol-1-yl]acetic acid) (purchased from the Albert Einstein College of Medicine of Yeshiva University), 0.4 mM CuSO_4_, 10 mM sodium ascorbate, and 10 mM aminoguanidine. Components of the reaction were added to CuAAC buffer (50 mM HEPES, 100 mM NaCl, 10 mM aminoguanidine, 0.1% DPC, pH 7.5) in the order of protein, tag, a premixed solution of BTTAA and CuSO_4_, and sodium ascorbate. Aminoguanidine was added to the buffer to prevent protein aggregation caused by byproducts from ascorbate oxidation. CuAAC reactions were terminated with 5 mM EDTA and allowed to stand in air for 30 minutes. The reactions were desalted using a Zeba spin desalting column (ThermoFisher Scientific) to remove excess C3-tag and concentrated by ultrafiltration using an Amicon centrifugal filter (30,000 MWCO).

### NMR sample preparation

An additional size exclusion step was used to remove any aggregation as a result of the CuAAC reactions using a Superdex® S200 Increase 10/300 GL column (Cytiva) equilibrated with NMR buffer (25 mM MES, 100 mM KCl, 0.15% DPC, pH 6.2). Pooled fractions were concentrated in an Amicon centrifugal filter (10,000 MWCO) and analyzed by SDS-PAGE to confirm the sample purity was greater than 95%. Final protein concentrations for each sample ranged from 0.5 – 0.7 mM in NMR buffer with 10% D_2_O.

### NMR spectroscopy

All NMR spectra were recorded on a 900 MHz Bruker Avance NMR spectrometer equipped with a cryoprobe at 313K. PCSs were measured from a single set of interleaved TROSY spectra using in phase/anti phase-heteronuclear single quantum coherence (IPAP-TROSY) experiments. All 2D ^1^H-^15^N TROSY-HSQC NMR spectra were processed with Bruker Topspin (version 3.2). PCSs were measured in the ^1^H and ^15^N dimensions of the ^1^H-^15^N TROSY-HSQC spectrum. PCSs for three different DsbB-pAzF mutants at positions Y97, V72 and F32, were collected in order to triangulate datasets and restrict the conformational search space for structural modeling in RosettaNMR.

### Magnetic susceptibility anisotropy (Δχ) tensor fits

Δχ tensors were determined using the experimental ^1^H and ^15^N PCSs of the backbone amides observed with Yb^3+^ by fitting to the first conformer in the DsbB NMR structural ensemble (PDB ID: 2K73^29^) using the program PyParaTools^47,48^. Initial estimations of the metal ion position(s) were given as the coordinates of the Cβ atom of the spin-label residue. Tensor fit quality was assessed by calculating the root mean-square deviation between experimental and back-calculated PCSs divided by the root-mean-square of the experimental PCSs to give a Q-factor^49^. Table 1 reports the Δχ tensors determined with the C3 tag loaded with Yb^3+^ at each pAzF position.

### Conversion of PCS and NOE data into Rosetta restraints

PCS data were evaluated by a Rosetta scoring method developed previously^40^. The PCS score is the sum of squared errors between the experimental and predicted PCS values for a given protein structure model. NOE data for DsbB were taken from BMRB accession 15966^29^ and used as atom pair distance restraints with a flat-bottom bounded penalty function. The applied penalty function grows quadratically outside the lower and upper bounds, and linearly at distances 0.5 Å larger than the upper bound. The upper bound was set to the experimental NOE distance plus an additional 1.5 Å padding, and the lower bound was set to 1.5 Å. In the fragment assembly calculation, NOE distances involving side chain protons were mapped onto the centroid (CEN) atom of an amino acid residue as described previously^50^.

The weights of the PCS and NOE restraint scores were adjusted such that the ranges of the restraint scores and Rosetta scores over all models were approximately equal. The weights were determined by first generating 10,000 models without NMR data and then rescoring them with PCS or NOE restraints. The weight was then calculated as 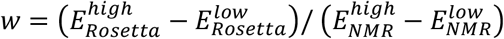. 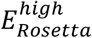 and 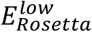 are the average of the highest and lowest 10% of the values of the Rosetta score, and 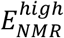 and 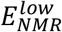 are the average of the highest and lowest 10% of the values of the PCS or NOE restraint score, respectively. For DsbB *de novo* structure prediction, *E_Rosetta_* represented the Rosetta score obtained with the score3 weight set.

### Rosetta structure calculations of DsbB

*De novo* structure prediction of DsbB was accomplished with Rosetta’s fragment assembly protocol^51^. PCS and NOE data were used in stages 1 to 4 of the fragment assembly protocol and for final model scoring and selection. The scoring function at each of the four assembly stages was supplemented with a score term for PCS restraints or NOE atom pair distance restraints combined with a weighting factor that was optimized against the total Rosetta score computed with the score3 scoring function. Backbone chemical shift data for DsbB were used for fragment selection and were taken from BMRB accession 15966^29^. The fragment search was carried out with the Rosetta3 fragment picker^52^ using secondary structure predictions created with PSIPRED^53^ and Jufo9D^54^. An additional fragment library was generated using secondary structure and φ/ψ angle predictions created with chemical shifts and the program TALOS+^55^. Homologous proteins of DsbB in the PDB with a PSI-BLAST E-value <0.5 were excluded from the fragment search. Structure calculations were performed with and without PCS data for DsbB, using either a fragment library created with PSIPRED and Jufo9D secondary structure information or a fragment library created with chemical shift data for DsbB. Additional structure calculations were performed with NOE restraints and combinations of PCSs and NOE restraints in order to compare the performance of the two types of NMR data. The different structure calculation experiments and number of NMR restraints are listed in Table S1. A total of 10,000 structure models of DsbB was generated for each of these structure calculations. Model accuracy was evaluated by calculating the Cα-atom RMSD over all residues and over residues in transmembrane segments (RMSD_TM) relative to the experimental reference structure of DsbB (PDB: 2K73)^29^.

