## Supporting Information for "Pseudocontact shift NMR data obtained from a non-canonical amino acid-linked lanthanide tag improves integral membrane protein structure prediction"

### Figure S1

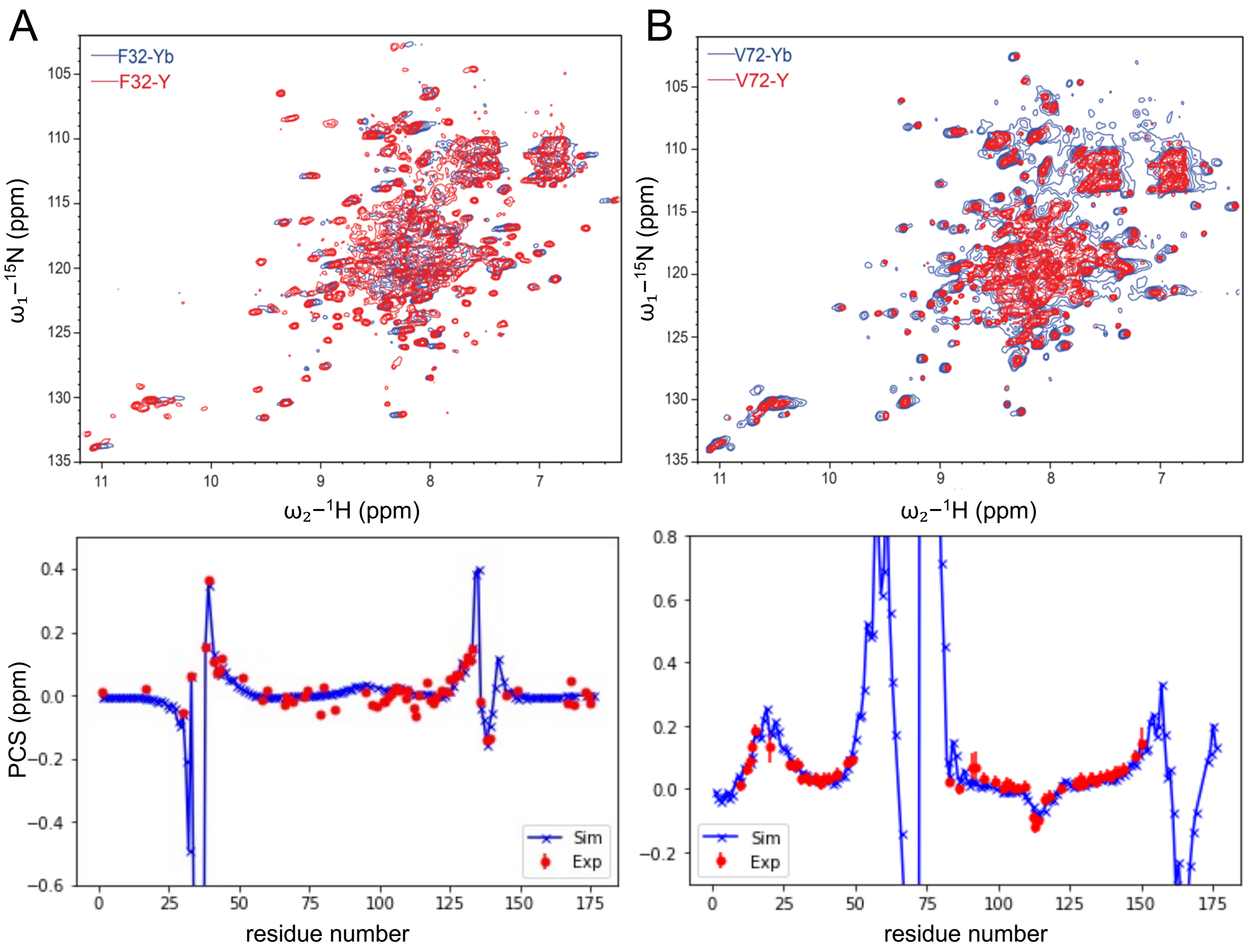


**Figure S1: $^{1}$H-$^{15}$N TROSY-HSQC NMR spectra for para- and diamagnetically-labeled DsbB at position F32pAzF and V72pAzF.** The $^{1}$H-$^{15}$N TROSY-HSQC spectrum for the paramagnetic C3-$Yb^{3+}$ sample (blue) is superimposed with the spectrum of the diamagnetic C3-$Y^{3+}$ sample (red) for DsbB labeled at positions A) F32pAzF and B) V72pAzF. The measured PCSs along the amino acid sequence for each position are shown below the respective NMR spectrum. NMR data was collected at 40°C and a proton resonance frequency of 900 MHz.

**Figure S2**

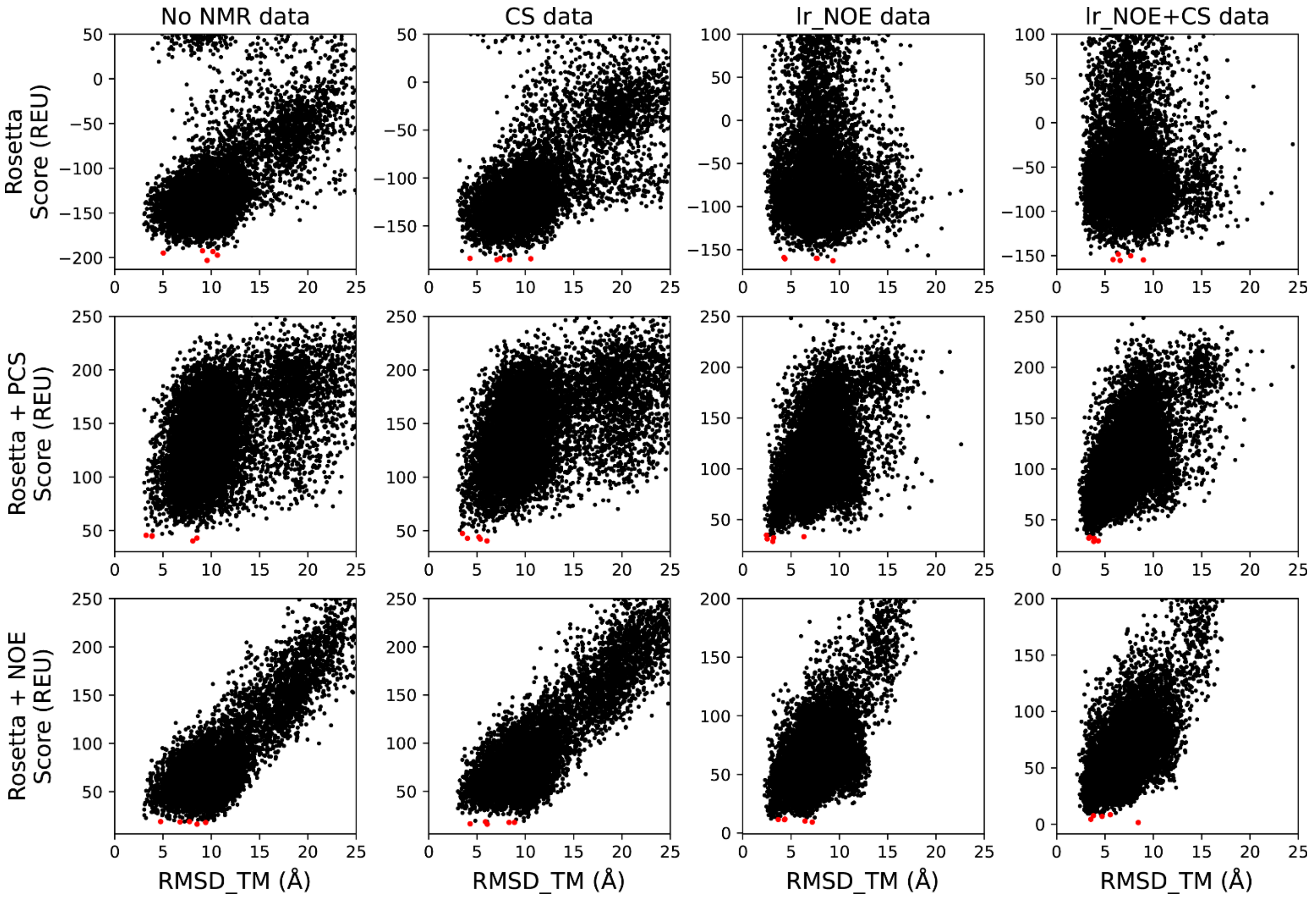


**Figure S2: Score-vs-RMSD plots of DsbB models predicted with Rosetta using chemical shifts and/or long-range NOE data.** Models were scored using the Rosetta score3 energy function (upper row) or a combination of the score3 energy and NMR restraint energy computed from either PCSs (middle row) or long-range NOEs (bottom row). The five lowest-scoring models are indicated by red dots.

**Figure S3**

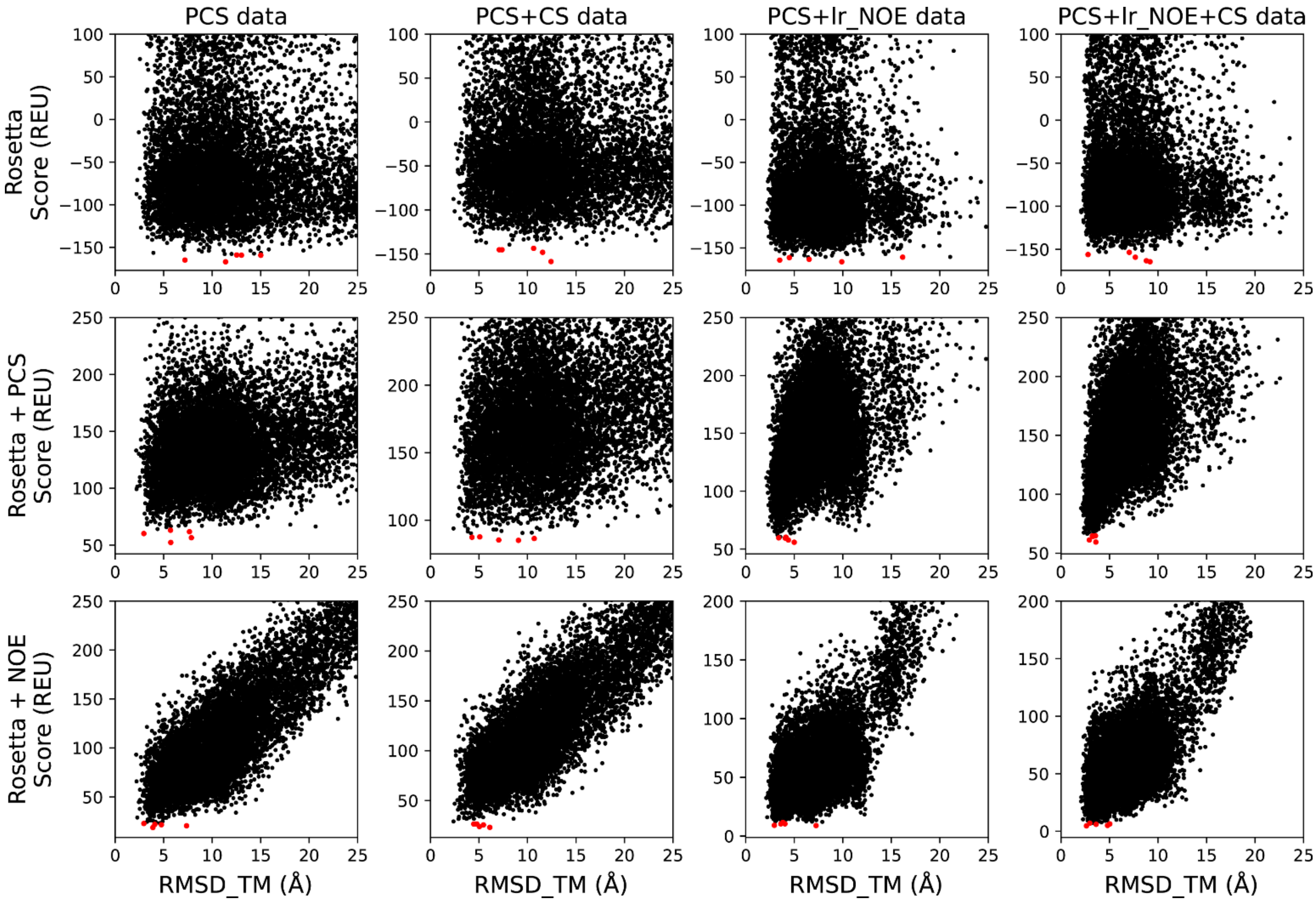


**Figure S3: Score-vs-RMSD plots of DsbB models predicted with Rosetta using PCS data alone or in combination with chemical shift data and/or long-range NOEs.** Models were scored using the Rosetta score3 energy function (upper row) or a combination of the score3 energy and NMR restraint energy computed from either PCSs (middle row) or long-range NOEs (bottom row). The five lowest-scoring models are indicated by red dots.

**Figure S4**

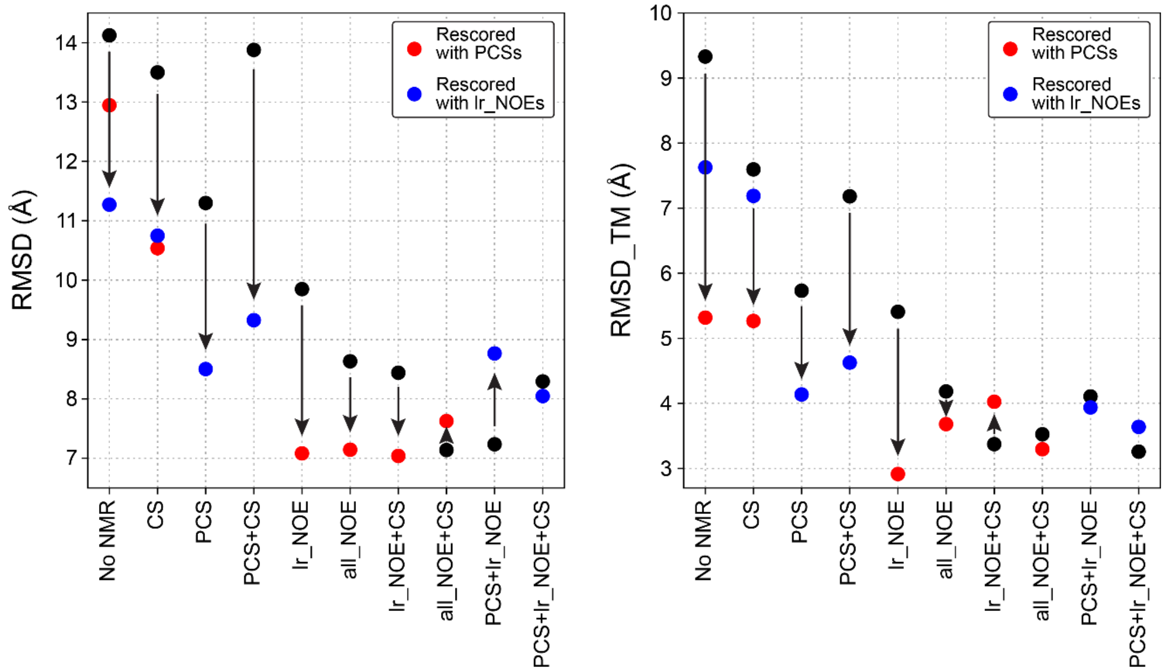


**Figure S4: Models with lower RMSD get selected when models are rescored with PCSs or long-range NOE data, which were not used in the *de novo* folding calculation.** The black data points indicate the Cα-RMSD (left) or Cα-RMSD_TM (right) values of DsbB models which were selected by scoring with the same types of NMR data used in the *de novo* folding simulation. Red and blue data points indicate the RMSD values of models selected after rescoring with either PCS or long-range NOE data. Black arrows indicate the direction of the RMSD change when models would be selected after rescoring.

**Table S1: Observed PCSs for each C3-Yb pAzF tagging site.**

| F32 | | | | V72 | | | | Y97 | | | |
|---|---|---|---|---|---|---|---|---|---|---|---|
| Resname | Residue | Atom | PCSs (ppm) | Resname | Residue | Atom | PCSs (ppm) | Resname | Residue | Atom | PCSs (ppm) |
| M | 1 | H1 | 0.0125 | Q | 10 | H | 0.0147 | M | 1 | H1 | 0.1712 |
| L | 17 | H | 0.0219 | R | 12 | H | 0.0625 | L | 2 | H | 0.1653 |
| L | 30 | H | -0.0559 | G | 13 | H | 0.0880 | R | 3 | H | 0.1397 |
| Q | 33 | H | 0.0610 | A | 14 | H | 0.1366 | L | 5 | H | 0.1174 |
| L | 38 | H | 0.1542 | W | 15 | H | 0.1864 | A | 8 | H | 0.0935 |
| K | 39 | H | 0.3641 | F | 20 | H | 0.1322 | S | 9 | H | 0.1338 |
| C | 41 | H | 0.1096 | L | 27 | H | 0.0767 | Q | 10 | H | 0.0827 |
| V | 42 | H | 0.0717 | A | 29 | H | 0.0712 | G | 11 | H | 0.0783 |
| L | 43 | H | 0.0826 | L | 30 | H | 0.0786 | R | 12 | H | 0.0535 |
| S | 44 | H | 0.1191 | W | 31 | H | 0.0332 | G | 13 | H | 0.0559 |
| L | 51 | H | 0.0582 | Q | 33 | H | 0.0384 | A | 14 | H | 0.0768 |
| A | 58 | H | -0.0139 | H | 34 | H | 0.0293 | W | 15 | H | 0.0776 |
| I | 60 | H | 0.0167 | M | 36 | H | 0.0325 | L | 17 | H | 0.0712 |
| K | 66 | H | -0.0306 | L | 37 | H | 0.0293 | M | 18 | H | 0.0750 |
| T | 67 | H | -0.0063 | L | 38 | H | 0.0186 | A | 19 | H | 0.0628 |
| L | 69 | H | -0.0207 | K | 39 | H | 0.0328 | F | 20 | H | 0.0524 |
| A | 73 | H | -0.0064 | C | 41 | H | 0.0340 | T | 21 | H | 0.0650 |
| M | 74 | H | 0.0184 | L | 43 | H | 0.0406 | A | 22 | H | 0.0569 |
| Y | 79 | H | -0.0581 | S | 44 | H | 0.0479 | L | 23 | H | 0.0197 |
| S | 80 | H | 0.0264 | E | 47 | H | 0.0818 | A | 24 | H | 0.0126 |
| G | 84 | H | -0.0425 | A | 49 | H | 0.0922 | A | 29 | H | -0.0050 |
| Q | 95 | H | 0.0094 | R | 83 | H | 0.0253 | L | 30 | H | 0.0000 |
| Y | 97 | H | -0.0284 | Q | 86 | H | 0.0001 | W | 31 | H | -0.0233 |
| S | 99 | H | -0.0328 | H | 91 | H | 0.0660 | Q | 33 | H | -0.0482 |
| F | 101 | H | -0.0205 | T | 92 | H | 0.0699 | H | 34 | H | -0.0312 |
| A | 102 | H | -0.0215 | Q | 95 | H | 0.0330 | M | 36 | H | -0.0498 |
| T | 103 | H | -0.0055 | S | 99 | H | 0.0220 | L | 37 | H | -0.0513 |
| S | 104 | H | 0.0075 | A | 102 | H | 0.0034 | K | 39 | H | -0.0237 |
| D | 105 | H | 0.0158 | T | 103 | H | 0.0181 | V | 42 | H | -0.0745 |
| F | 106 | H | 0.0281 | S | 104 | H | 0.0113 | L | 43 | H | -0.1276 |
| M | 107 | H | 0.0177 | D | 105 | H | 0.0039 | S | 44 | H | -0.1032 |
| R | 109 | H | -0.0105 | M | 107 | H | 0.0000 | Y | 46 | H | -0.0207 |
| F | 110 | H | 0.0173 | R | 109 | H | 0.0074 | E | 47 | H | -0.0721 |
| E | 112 | H | -0.0401 | E | 112 | H | -0.0901 | R | 48 | H | -0.0449 |
| W | 113 | H | -0.0638 | W | 113 | H | -0.1212 | A | 49 | H | -0.0538 |
| L | 114 | H | 0.0020 | L | 114 | H | -0.0981 | A | 50 | H | -0.0572 |
| D | 117 | H | 0.0412 | L | 116 | H | -0.0327 | L | 51 | H | 0.0510 |
| K | 118 | H | -0.0143 | K | 118 | H | -0.0243 | G | 53 | H | 0.0091 |
| W | 119 | H | -0.0139 | Q | 122 | H | 0.0034 | V | 54 | H | 0.0197 |
| V | 120 | H | -0.0036 | S | 127 | H | 0.0192 | G | 56 | H | 0.0291 |
| Q | 122 | H | 0.0148 | G | 128 | H | 0.0147 | A | 57 | H | 0.0538 |
| V | 124 | H | 0.0095 | D | 129 | H | 0.0327 | A | 58 | H | 0.0578 |

| | | | | | | | | | | | |
|---|---|---|---|---|---|---|---|---|---|---|---|
| F | 125 | H | 0.0512 | C | 130 | H | 0.0174 | L | 59 | H | 0.0544 |
| A | 126 | H | 0.0513 | E | 132 | H | 0.0327 | G | 61 | H | 0.0465 |
| S | 127 | H | 0.0644 | R | 133 | H | 0.0324 | A | 62 | H | 0.0513 |
| G | 128 | H | 0.0670 | Q | 134 | H | 0.0251 | I | 63 | H | 0.0403 |
| C | 130 | H | 0.0984 | D | 136 | H | 0.0400 | A | 64 | H | 0.0511 |
| A | 131 | H | 0.1205 | L | 138 | H | 0.0440 | T | 67 | H | 0.0561 |
| E | 132 | H | 0.1101 | G | 139 | H | 0.0366 | L | 69 | H | 0.0270 |
| R | 133 | H | 0.1462 | E | 141 | H | 0.0513 | R | 70 | H | 0.0607 |
| D | 136 | H | -0.0202 | M | 142 | H | 0.0547 | Y | 71 | H | 0.0705 |
| L | 138 | H | -0.1404 | Q | 144 | H | 0.0626 | V | 72 | H | 0.0521 |
| G | 139 | H | -0.1372 | G | 148 | H | 0.1026 | A | 73 | H | 0.0630 |
| W | 145 | H | -0.0003 | F | 150 | H | 0.1466 | M | 74 | H | 0.0861 |
| I | 149 | H | 0.0185 | | | | | V | 75 | H | 0.0845 |
| K | 167 | H | -0.0244 | | | | | W | 77 | H | 0.0939 |
| A | 168 | H | 0.0444 | | | | | Y | 79 | H | 0.0675 |
| K | 169 | H | -0.0309 | | | | | S | 80 | H | 0.0184 |
| L | 173 | H | 0.0090 | | | | | R | 83 | H | -0.0187 |
| F | 174 | H | 0.0002 | | | | | G | 84 | H | 0.0050 |
| G | 175 | H | -0.0252 | | | | | V | 85 | H | -0.0025 |
| W | 119 | HE | -0.0227 | | | | | T | 92 | H | -0.5508 |
| | | | | | | | | L | 94 | H | -0.5379 |
| | | | | | | | | A | 102 | H | -0.0264 |
| | | | | | | | | T | 103 | H | -0.0459 |
| | | | | | | | | D | 105 | H | -0.0450 |
| | | | | | | | | F | 106 | H | -0.0484 |
| | | | | | | | | M | 107 | H | -0.0498 |
| | | | | | | | | V | 108 | H | -0.0805 |
| | | | | | | | | R | 109 | H | -0.0666 |
| | | | | | | | | F | 110 | H | -0.0502 |
| | | | | | | | | E | 112 | H | -0.0346 |
| | | | | | | | | W | 113 | H | -0.0391 |
| | | | | | | | | L | 114 | H | -0.0219 |
| | | | | | | | | L | 116 | H | -0.0119 |
| | | | | | | | | D | 117 | H | -0.0121 |
| | | | | | | | | K | 118 | H | -0.0311 |
| | | | | | | | | W | 119 | H | -0.0300 |
| | | | | | | | | V | 120 | H | -0.0130 |
| | | | | | | | | Q | 122 | H | -0.0136 |
| | | | | | | | | A | 126 | H | -0.0821 |
| | | | | | | | | S | 127 | H | -0.1092 |
| | | | | | | | | G | 128 | H | -0.0963 |
| | | | | | | | | D | 129 | H | -0.0507 |
| | | | | | | | | C | 130 | H | -0.0989 |
| | | | | | | | | E | 132 | H | -0.2435 |
| | | | | | | | | R | 133 | H | -0.2481 |
| | | | | | | | | Q | 134 | H | -0.2164 |
| | | | | | | | | W | 135 | H | -0.2022 |

| | | | |
|---|---|---|---|
| D | 136 | H | -0.3618 |
| L | 138 | H | -0.2394 |
| G | 139 | H | -0.4013 |
| I | 162 | H | 0.1346 |
| S | 163 | H | 0.1441 |
| Q | 164 | H | 0.1119 |
| F | 166 | H | 0.0702 |
| A | 168 | H | 0.0688 |
| K | 169 | H | 0.0518 |
| F | 174 | H | 0.0345 |
| G | 175 | H | 0.0210 |
| W | 15 | HE1 | 0.0800 |
| W | 31 | HE1 | -0.0157 |
| W | 119 | HE1 | -0.0166 |
| W | 135 | HE1 | -0.1039 |

**Table S2: Overview of Rosetta structure calculations for DsbB and used NMR data.**

| | NMR data type | No. PCSs | No. NOEs | Data source |
|---|---|---|---|---|
| 1 [a] | no PCS or NOE | - | - | - |
| 2 | PCS | 60 / 54 / 99 [b] | | this study |
| 3 | lr_NOE [c] | | 44 | BMRB: 15966 |
| 4 | all_NOE [d] | | 500 | BMRB: 15966 |
| 5 | PCS + lr_NOE | 60 / 54 / 99 | 44 | this study and BMRB: 15966 |

[a] Calculations 1 - 5 were performed with two different fragment libraries, one calculated with PSIPRED and Jufo9D secondary structure predictions, and another calculated with chemical shift-derived secondary structure predictions. Backbone chemical shifts of DsbB were taken from BMRB: 15966.

[b] Number of $H^N$ PCSs determined at positions F32, V72, and Y97.

[c] A proton pair was considered a long-range NOE if the sequence separation between residues was ≥5.

[d] Includes number of short-range and long-range NOEs
